# Unsupervised idealization of ion channel recordings by Minimum Description Length: Application to human PIEZO1-channels

**DOI:** 10.1101/106187

**Authors:** Radhakrishnan Gnanasambandam, Morten Schak Nielsen, Christopher Nicolai, Frederick Sachs, Johannes Pauli Hofgaard, Jakob Kisbye Dreyer

## Abstract

Researchers can investigate the mechanistic and molecular basis of many physiological phenomena in cells by analyzing the fundamental properties of single ion channels. These analyses entail recording single channel currents and measuring current amplitudes and transition rates between conductance states. Since most electrophysiological recordings contain noise, the data analysis can proceed by idealizing the recordings to isolate the true currents from the noise. This de-noising can be accomplished with threshold crossing algorithms and Hidden Markov Models, but such procedures generally depend on inputs and supervision by the user, thus requiring some prior knowledge of underlying processes. Channels with unknown gating and/or functional sub-states and the presence in the recording of currents from uncorrelated background channels present substantial challenges to unsupervised analyses.

Here we describe and characterize an idealization algorithm based on Rissanen’s Minimum Description Length (MDL) Principle. This method uses minimal assumptions and idealizes ion channel recordings without requiring a detailed user input or a *priori* assumptions about channel conductance and kinetics.. Furthermore, we demonstrate that correlation analysis of conductance steps can resolve properties of single ion channels in recordings contaminated by signals from multiple channels. We first validated our methods on simulated data defined with a range of different signal-to-noise levels, and then showed that our algorithm can recover channel currents and their substates from recordings with multiple channels, even under conditions of high noise. We then tested the MDL algorithm on real experimental data from human PIEZO1 channels and found that our method revealed the presence of substates with alternate conductances.

## Introduction

The analysis of discrete events in single ion channel data has been a powerful tool in electrophysiology research since the pioneering work of Erwin Neher and Bert Sakmann, recognized with the 1991 Nobel Prize in Medicine. However, these analyses usually entail data modeling methods that rely on user-defined inputs, filters, event detection thresholds or subjective criteria for event detection. These conditions present difficulties, particularly when analyzing time series composed of currents from multiple ion channels, or in cases when a channel can make a transition between one or more sub-conductance states.

Given these considerations, approaches to the analysis of ion channel currents often involve one of two paradigms. In one approach, Hidden Markov Models (HMMs) are used to analyze recordings containing signals arising from one or more ion channels (Qin, 2007). Here, we assume that the ion channels generate the observed currents by jumping between metastable conductance states. Consequently, HMM analysis is most suitable when different conductance states of channels can be estimated a *priori*; so long as these estimates are valid, the HMM algorithm provides maximal information about the conductance transitions and their kinetics.

HMM is not always applicable, precisely because it requires a *priori* knowledge of likely kinetic models, which can be especially difficult to predict if currents from several channels are present in the recording. While the HMM-based algorithms can provide useful quantitative analyses, real laboratory data often fail to satisfy the inherent assumptions of the model. Chief among these assumptions or conditions is that the system should be in a steady-state for the duration of the recording. However, the channel kinetics and ion flux can vary with time, for example the membrane resting potential may change during the recording (Suchyna et al., 2009; Gottlieb et al., 2012). An alternative to the HMM approach is to idealize the measured input sequence using an event detection algorithm that is independent of the kinetic model (Colquhoun and Sigworth, 1995; Carter et al., 2008; Parsons and Huizinga, 2013), The measured current is thus modeled as a series of events without imposition of a particular kinetic scheme, and can in principle account for unknown sub-states in the recording. Methods that analyze the filtered time course of conductance transitions enable highly accurate determination of transition points (Colquhoun and Sakmann, 1985). However, even the most advanced methods have previously required user-defined inputs such as event trigger thresholds and the selection of some particular low-pass filter cut-off frequency.

As an alternate to these methods, we now test Rissanen’s Minimum Description Length Principle (MDL) to provide an idealization algorithm based on “structural breakpoint detection” (Rissanen, 1978;Lee, 2001;Davis et al., 2006;Killick et al., 2012). Our aim is to provide a fast and unbiased idealization of single channel time-series without requiring any user-dependent a *priori* model inputs. We assume that the current can be modeled stepwise as a sequence of segments of constant amplitude, separated by abrupt (instantaneous) transitions to some new current amplitude. The algorithm identifies the location of the transitions between segments, and calculates the mean current prevailing in each segment. Furthermore, the MDL algorithm performs the calculation of transition locations and amplitudes without a *priori* assumptions of step-size or transition kinetics, and the current amplitude in each segment is independent of other amplitudes occurring in the recording. To provide a post hoc analysis of the output data, we developed a correlation analysis in which the conditional probability of observing pairwise adjacent steps is used to infer the number of discrete states. Finally, we tested the MDL algorithm on real experimental data from human PIEZO1 channels and found that our method revealed the presence of substates with alternate conductances.

## Methods

### Cell culture and Electrophysiology

We analyzed current records from the mechanically gated channel PIEZO1 in transformed HEK cells maintained in an incubator at 37°C and 5% CO_2_. These cells had been transfected with 200-500 ng of hPIEZO1 cDNA one-two days prior to performing cell-attached patch-clamp recordings at room temperature. The resting membrane potential of cells was maintained close to 0 mV by using a high potassium bath solution containing 150 mM KCl, 10 mM HEPES, and 1 mM MgCl_2_ and CaCl_2_, adjusted to pH 7.4. The pipette solution contained 150 mM KCl, 80 mM TEA, and 10 mM HEPES at pH 7.4. During the recordings, the patch membrane potential was randomly stepped to voltages within the range of -100 mV to +100 mV and pressure steps were applied to the pipette using a high-speed pressure clamp (ALA Scientific) (Figure 6A and B). Data sampling was at 10 kHz, with Bessel filtration at 2 kHz.

### Mathematical Analysis

The basic assumption in our method is that the opening and closing of an ion channel gives rise to instantaneous changes in membrane current, which are superimposed on a noisy background of the recording. Accordingly, we modeled the time series of currents as a sequence of steps corresponding to opening and closing of channels and additive Gaussian noise (ignoring for the present the possibility of state-dependent noise). We define time points at which the conductance jumps occur as break points, and the difference in mean amplitude on each side of the break point as step height (or step amplitude).

Consider a data set denoted as ***x** = (x_1_, x_2_, …, x_N_)* consisting of *N* data points. Here, we define the code length *L* as the length of the message containing the relevant information in the data. The code length, *L*, need not be constant, but varies depending on how the information in ***x*** is represented. More effective coding schemes generally have low description length. In particular, if ***x*** is well described by a particular model, it may be more efficient to first encode the model parameters, and then encode any deviations from the model (Rissanen, 1978; Lee, 2001; Davis et al., 2006). MDL serves to rank the fitness of models of a data set by identifying the particular model producing the shortest total description length for the observations; in other words, the task is to select the model providing an optimal tradeoff between model complexity and fitting of the data.

In order to apply MDL to an ion channel recording, we first need to calculate the lower bound of the description length of the observed time series, ***x***. Considering a time series segment ***x** = (x_1_, x_2_, …, x_N_)* with mean value *μ*, the residual sum of squares (RSS) is defined as

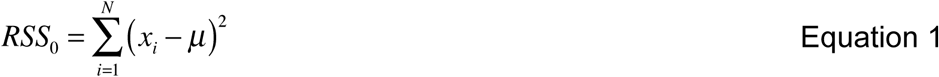

We now consider the minimum description length for encoding the data sequence (Rissanen, 1978; Lee, 2001). A set of *N* data points with a Gaussian distribution can be most efficiently encoded as the mean value and a list of deviations from the mean. Thus, the minimum description length of a segment described by a single mean value is given as

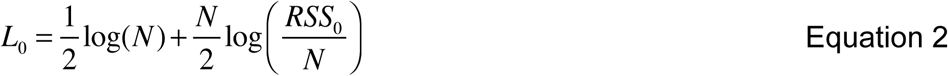

In Equation 2 the first term is the description length of the mean value; the minimum number of bits required to encode the mean value depends on the precision with which it is determined, and that precision is determined by *N*, the number of data points. The second term of the equation is the description length of the *N* deviations from the mean. By convention of information theory, base 2 logarithms (log_2_) are usually used to calculate code length in units of bits. However, for the application at hand, we are interested in comparing relative changes in code length, and therefore prefer to use the natural logarithm, the particular base being irrelevant.

We next consider the same sequence of data, but divided into daughter segments at *k* locations defined by *2 ≤ n_i_ ≤ N-1*, and where *n_i_ < n_i+1_*. We then define the residual sum of squares of the divided segment (RSS_*k*_) as

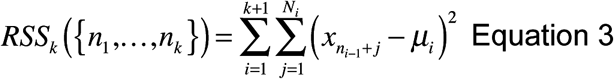

where *n_i_* is the *i*’th division point and *μ_i_* is the mean value of the *N_i_* data points between *n_i-1_* and *n_i_*. As above, we then calculate the minimum description length of the divided segment as (Lee, 2001),

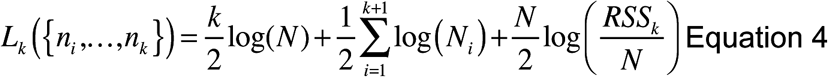

Here the first term describes the code length for the location in the data string of the defined break points, i.e. the points at which the segment is divided into *k+1* daughter segments. The second term describes the code length of the mean value of each segment, and the third term is the code length for the residuals. According to the Minimal Description Length Principle, a divided segment can be considered a better model of the data only if *L_k_ < L_0_*. In our implementation, we applied Equation 4 with *k=1* or *k=2*.

### Step detection algorithm

Having come this far, the task is now to identify a set of breakpoints minimizing the description length of a given dataset. This is inherently a complex multidimensional optimization problem. For typical data the number of possible segmentations is enormous and a full search for all possible breakpoints not feasible. We therefore search for breakpoints iteratively using a modification of the Binary Segmentation process (Scott and Knott, 1974; Kalafut and Visscher, 2008). In brief, we try to locate 1 or 2 breakpoints in the full segment and, if successful, we repeat the search on each sub-segment. We first try to locate a single break point in the time series. If this putative breakpoint fails to reach the MDL criterion, we try to locate two break points. Thus, our search is essentially a *tertiary segmentation* process with an initial binary search step. Using numerical tests, we show below that for the binary search, the probability of detecting channel-like events depends on the recording length, but that this undesirable property is circumvented with the tertiary search.

The algorithm proceeds in the following steps:

1. Calculate *L_0_* for the segment.
2. Search for the optimal location for a single break point (*k=1*). In other words, find *n\** as the *n* where *RSS_1_* is minimized. Thus, we define *n\** as

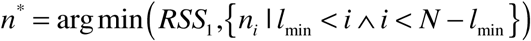

where *l*_min_ is a cutoff threshold for the smallest segment allowed. Unless otherwise stated we use *l*_min_ = 3.
3. Apply MDL to test whether division at the optimal breakpoint, *n\**, is indeed a better model than for the entire undivided segment:

a. Calculate *L_1_(n\**) using Equation 4
b. If *L_1_(n*) < L_0_* the segment is divided at *n\** and the procedure is repeated from step 1 for each of the two daughter segments. Thus, if the single breakpoint is accepted in step 3b, the algorithm is repeated from step 1. Otherwise, we proceed by evoking the tertiary search noted above, thus trying to locate two breakpoints in the segment:
4. Find the pair {*m_1_*, m_2_*}* for which *RSS_2_* is minimized.
5. Apply MDL to test if a model with divisions at the two breakpoints, {*m_1_*, m_2_*}*, provides a better model than the undivided segment:

a. Calculate *L_2_({m_1_*, m_2_\**}) using Equation 4
b. Accept {*m_1_*, m_2_\**} as break points if *L_2_({m_1_*, m_2_*}) < L_0_*
6. If the breakpoints are accepted, we repeat the procedure from step 1 onward for each daughter segment.

The algorithm proceeds until no sub-segment is further divisible (into two or three parts). Ultimately, the algorithm produces a list of breakpoints and the mean value of the segments follows easily.

The number of calculations using the tertiary search in step 4 scales with *N^2^*. However, in data containing many breaks, the binary search, i.e. steps 1-3.b of the search algorithm, is often by itself sufficient to decompose the sequence into shorter segments. This reduces the total computing time because the computationally demanding tertiary search is evoked on shorter sub-segments later in the search (see **Figure 3**).

We tested a number of alternate criteria for the detection of break points. Similar results were obtained when putative break points were tested using the *Bayes Inference Criterion* (BIC) (Schwarz, 1978), although a minimum segment length (*l*_min_=10) was required to avoid false segmentation of small segments. The Akaike Information Criterion (AIC) (Akaike, 1973) allowed many false steps, and was thus considered inappropriate in the present application (data for BIC and AIC are not shown). Similar observations have been reported by Kalafut and Visscher (2008), who used BIC in their event detection algorithm. We note that the alternate criteria BIC and AIC both require *a priori* knowledge of the variance of the noise. While this is estimable from a segment of the time series in which no steps/transitions were apparent by visual inspection, such a manual approach introduces bias favoring higher baseline noise models. However, the MDL method does not require this potentially subjective operation, and has superior performance in terms of low number of false positives.

The step detection algorithm with MDL was implemented in MATLAB and is available on MATLAB central file exchange (Dreyer, 2016). It is also available as a plugin for ion-channel analysis software QuB (https://www.qub.buffalo.edu/download/).

### Computational Complexity

The RSS of *N* points was computed with *O(N)* time complexity, using a well-known incremental one-pass method (Chan et al., 1983).

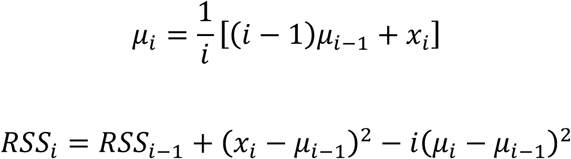

We derived the decrimental form, which removes a point from the distribution by solving for *μ_i-1_* and *RSS_i-1_*.

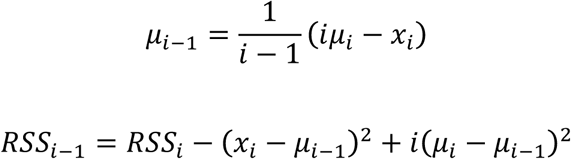

With these methods, the optimal break point, *n\**, was found with *O(N)* complexity. We initialized the left-hand distribution *D_L_* with the first *l_min_* points, and the right-hand distribution *D_R_* with all *N* points. The *RSS_2_* of the first break point candidate is then the sum of the left- and the right-hand incremental *RSS* values. We computed the *RSS_2_* of each subsequent candidate by adding a point to *D_L_* and removing it from *D_R_*. The tertiary search for the pair {*m_1_*, m_2_\**} was implemented by evaluating, for all possible m_1_, the optimal binary division of points *m_1…N_*, thus meeting the criterion for *O(N^2^)* complexity.

The overall complexity of the algorithm depends on the scale of the input data and the order in which intervals are segmented. Considering for example the binary segmentation of an interval sequence ABCDEFGH (i.e.*N_I_*=8 intervals), in the best possible case this might be decomposed first as ABCD, EFGH, then AB, CD, EF, GH, and finally A, B, C, D, E, F, G, H. The tree’s height, or in other words the number of generations, is then *g=log_2_(N_I_)*. In the worst case, the sequence might be first decomposed into A, BCDEFGH, then A, B, CDEFGH, and so on, and *g = N_I_-1*, or *O(N_I_)*. Since each generation involves a linear pass through the data, the intervals will be fully split with *O(gN)* complexity. If signal-to-noise ratio (SNR) is low, intervals are found by the tertiary search, which increases the complexity to *O(gN^2^)*. Thus, in many applications, the SNR is a critical factor influencing computational burden; when SNR is high, most steps are found using a single point search and computation time grows like *O(gN)*, but when SNR is low, double point search dominates, such that computation time grows like *O(gN^2^)*.

We tested the computation time of the MDL algorithm and compared with timing of HMM based segmentation using ‘viterbi_path’, a MATLAB implementation of the Viterbi algorithm coded by Kevin Murphy (Murphy, 1998). We performed the test on a Macbook pro (2.3 GHz Intel Core i7) using the builtin MATLAB timer functions ‘tic’ and ‘toc’. We took care to ensure that the process used more than 99% of resources in a single CPU core.

### Coherence spectra

We determined the coherence of the idealized output of the sequence with the input sequence as a function of noise level (**Figure 2**). At each defined SNR, we generated five data sequences, each consisting of 131072 data points, using a single Markov process. We average the coherence spectra in sections of 1024 points with 50% overlap and Hamming window using the MATLAB function ‘mscohere’.

### Correlation between neighboring events

The outputs of our algorithm are a list of breakpoints indicating the locations where the current is found to change abruptly, and the step amplitudes at these locations. Since the breakpoints and step amplitudes are determined independently, one therefore needs to undertake post-processing in order to analyze properties of single channels, in particular when there are transitions between states of various condutance.

We define the step amplitude *s_i_* is defined as *μ_i+1_ - μ_i_*. The precision of *s_i_* compared to the true value is given by

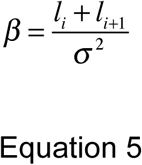

where *l_i+1_* and *l_i_* is the length of each of the adjacent segments and σ^2^ is the variance of the noise.

Consider a list of *M* step amplitudes, *s_1_, s_2_, …, s_M_* determined from an ion channel recording. We denote by *P(x)* the probability of observing a step of size *x*, and calculate this probablity as

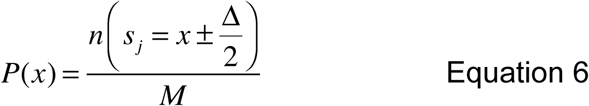

Here, the numerator indicates the number of steps with amplitude x and within a bin-width Δ, and the denominator is the total number of observed steps.

Our aim is to determine how often steps of amplitude *x* were immediately followed by steps of amplitude *y* in the recording. To this end, we therefore first define *P_xy_* as the joint probability of observing a step of amplitude *x* being immediately followed by a step of amplitude *y*:

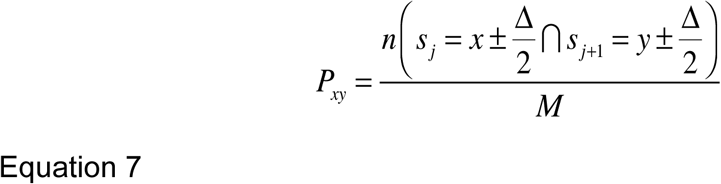

Here the numerator indicates the number of steps with amplitude x±Δ/2 followed by steps of amplitude y±Δ/2. The correlation between subsequent neighboring steps was then calculated as

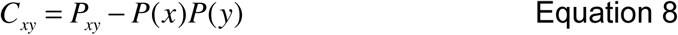

The sign of C indicates if particular transitions are over- or underrepresented relative to an independent process, and may thus provide physiologically relevant details of the underlying step-generating mechanism. In particular, *C_xy_* > 0 if the observation “*step of size x is followed by step of size y*” happens more frequently than if the observed sequence was purely random and *C_xy_* < 0 if it is less frequent. We used the MATLAB function ‘hist3’ to generate joint probability histograms, selecting a bin width Δ in the range 5-10% of the typical maximal amplitude. The resultant joint probability distributions, *P_xy_* and *C_xy_*, were smoothed using a 2D Gaussian kernel with a half-maximum width of 1 bin-width. If either *P(x)*=0 or *P(y)=0* (that is to say, if no events occurred with amplitudes x±Δ/2 or y±Δ/2), the correlation is undefined. In the analysis below we consider regions where the bin counts in the joint probability histogram ≤ 1 as undefined and these regions are indicated by white in the plots.

### Generation of simulated data

We used simulated data for evaluating the basic properties of our step detection method (**Figure 1** – **Figure 5**). As test data for quantifying the reliability of our detection mechanism, we simulated a simple two-state ion channel with the transition probability between conduction states being 1/100 per time step (**Figure 1**). The kinetics and noise in the test data were deliberately defined so as to challenge the limits of our detection method. Test data for quantifying the false positive detections consisted of homogeneous sequences of random numbers with a Gaussian distribution. This process was also used for coherence spectra (**Figure 2**) and timing measurements (**Figure 3**)

**Figure 1:**
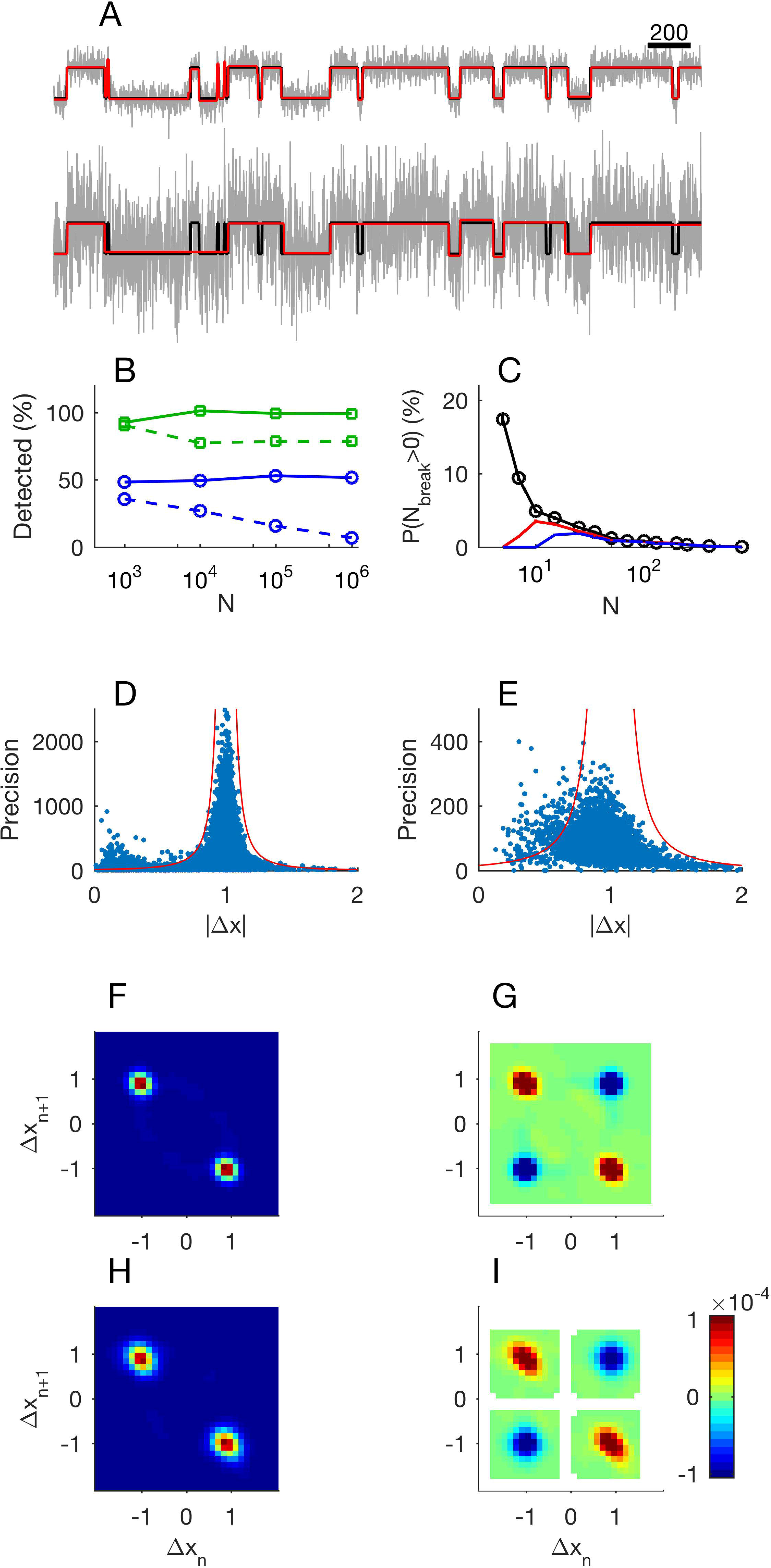
Evaluation of the step detection method applied to synthetic data. Panel A: 1000 data points of unit steps (upper, SNR = 3.3; lower, SNR = 1). Input data are shown in black, MDL-idealized data are shown in green, and true states are shown in red (note that the green and red lines often coincide). Panel B: The tertiary search method provides a constant frequency of event detection regardless of recording segment length. Markers show the number of detected steps relative to the number of known steps as a function of *N*, the number of data points in the record. Squares (green).: SNR=3.3 Circles (blue): SNR = 1. Solid lines show results from the tertiary search, and dashed lines show the results from the binary search. Panel C: Analysis of probability of false events. Each data point shows mean number of false events from 5000 sequences of uniform random numbers of length *N*. Black: *l*_min_ = 1, red: *l*_min_ = 3, and blue: *l*_min_ = 5. Panel D and E: Analysis of precision in step estimates at SNR 3.3 and 1 respectively. Dots show precision of jump estimate as function of observed absolute jump. We calculated the precision using the length of the adjacent segments according to Equation 5. Red lines indicate 4 standard deviations from the target value at different levels of precision. Panel F: Histogram of neighboring events at SNR 3.3. The *x*-axis shows *Δx* of event *n* and the *y*-axis shows *Δx* of the following event, *n*+*1*. Panel G: Correlation of events at SNR 3.3. Detected events display same correlations of neighboring events as a simple channel. Red indicates neighboring events occurring more frequently than random. Blue indicates neighboring events occurring less frequently than random. White areas indicate undefined correlation. Panel H: Histogram of neighboring events at SNR 1. Panel I: Correlation of events at SNR 1.

**Figure 2:**
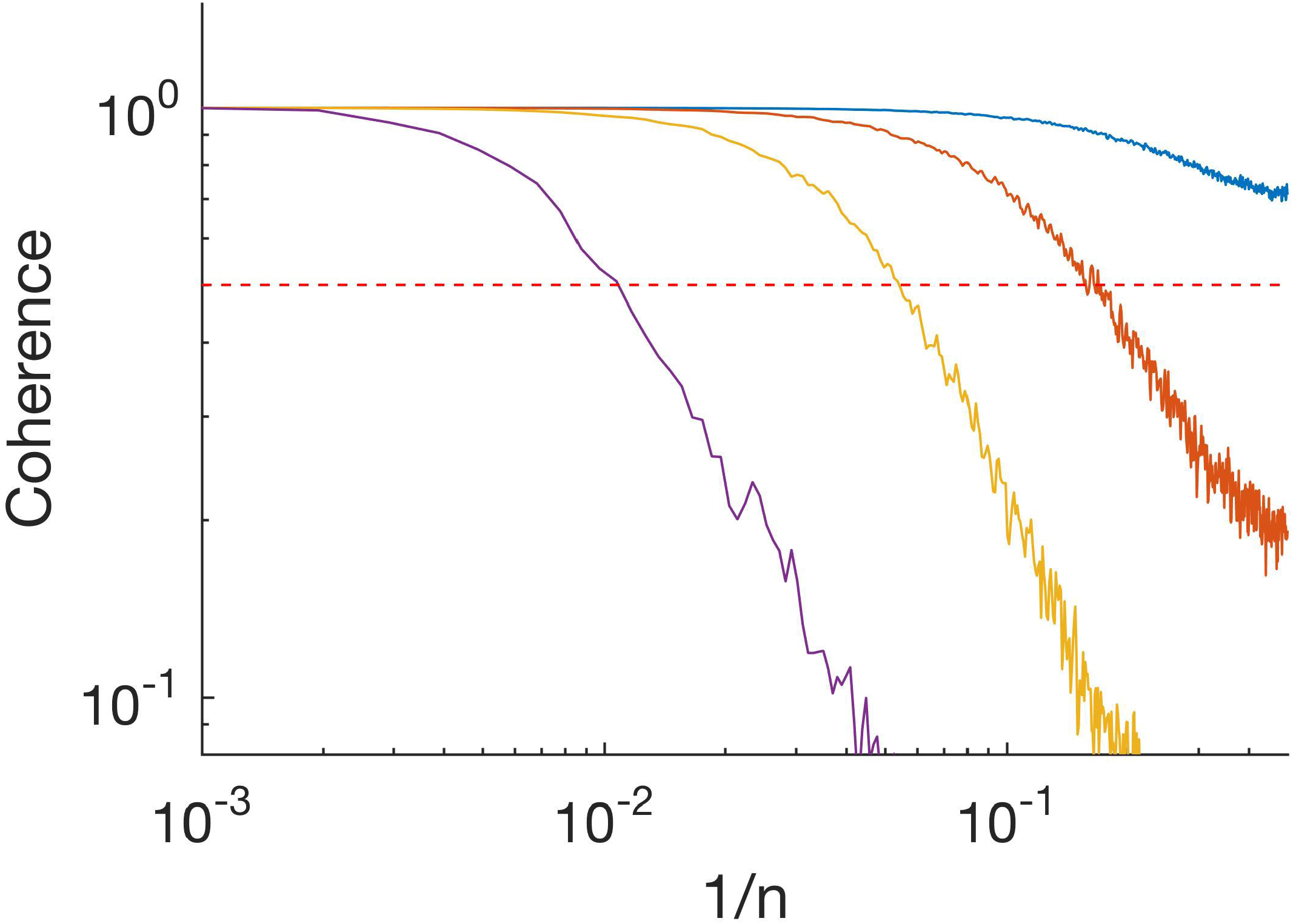
Absolute value of coherence spectrum between input time series and idealized time series. Purple, SNR = 1; yellow, SNR = 3.3; orange, SNR = 10; and blue, SNR = 33. Frequency is plotted as *1/n* where *n* is the length of the segment. Horizontal red dashed line indicates 0.5.

**Figure 3:**
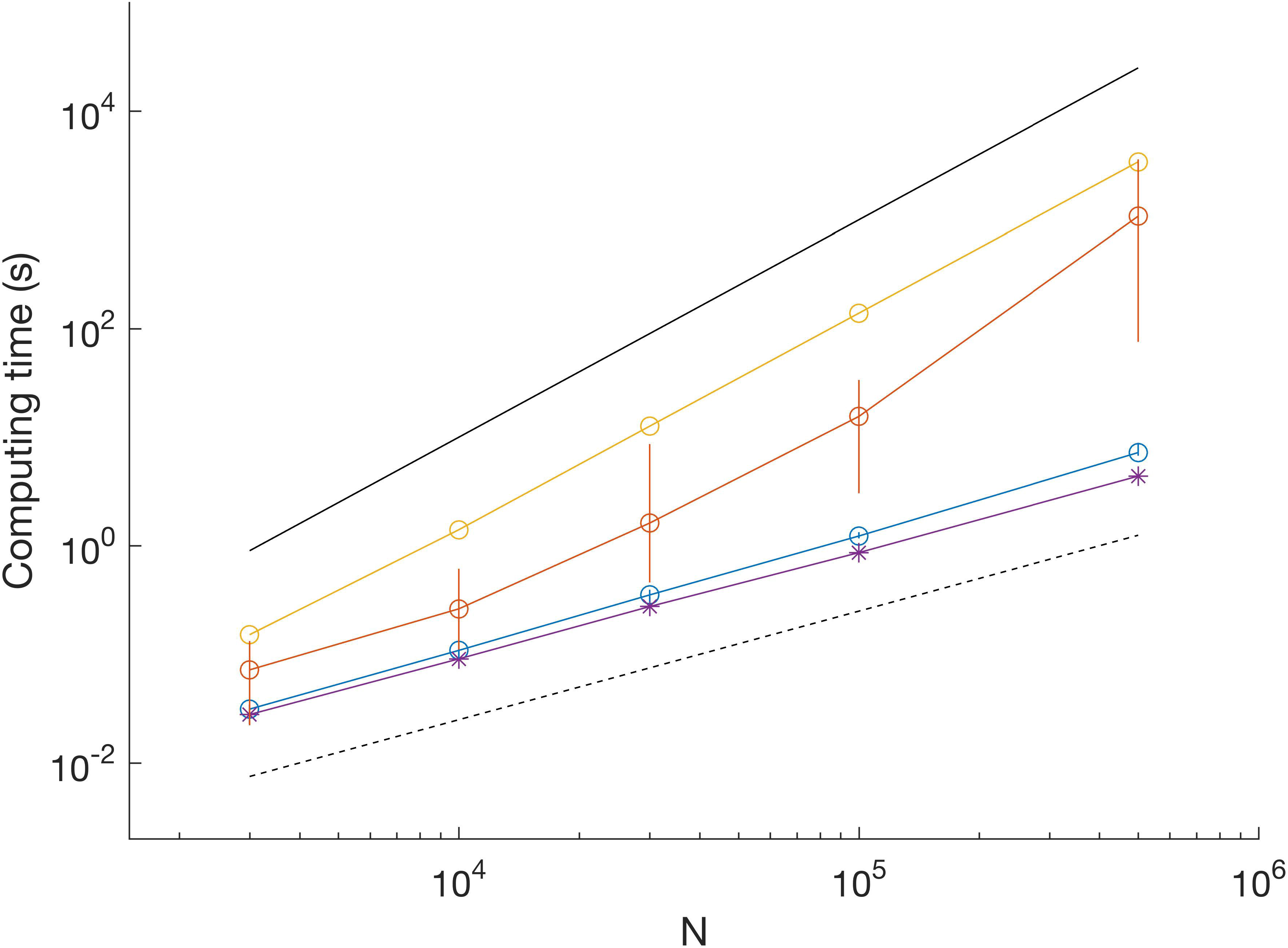
Comparison of computation time for different datasets. SNR = 3.3; blue dots show mean and vertical lines show range between minimum and maximum from 10 iterations. SNR = 1; orange dots show mean and vertical lines show range from 10 iterations. SNR = 0 (uniform random data), yellow, single iteration. Example of scaling by Viterbi algorithm (SNR = 3.3); purple asterisk, single iteration. Black lines indicate *O(N)* (dashed) and *O(N^2^)* scaling (solid).

**Figure 4:**
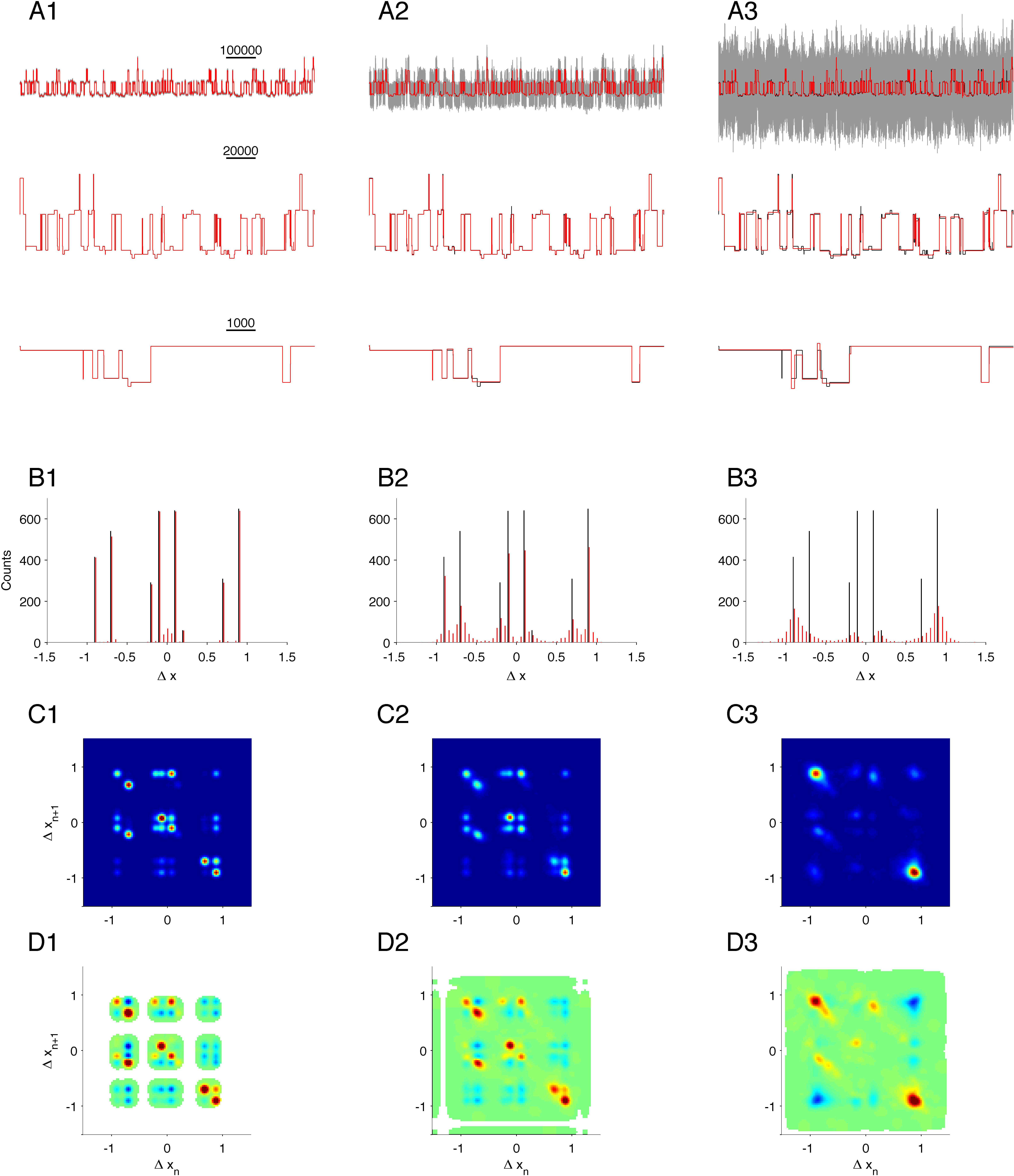
Event detection using different algorithms on simple simulated data. Panel A: Top; input data (gray) and Markov states (black). Blue states detected using threshold crossing algorithm applied on low-pass filtered data (low-pass filtered raw data are shown in black in the lower traces of A); green, states detected using Viterbi-algorithm as implemented in QuB. Red: states detected using MDL-algorithm. Panel B: Distribution of step-lengths. Black, step lengths of input; solid blue, threshold crossing on low-pass filtered data (*F_Ny_ /40*); green, Viterbi; red, MDL; cyan, threshold crossing on light low-pass filtered data (*F_Ny_ /20*). Panel C: Close-up view of distribution of step lengths. Cyan, threshold crossing on heavily low-pass filtered data (*F_Ny_ /80*). Other colors as in Panel B. Panel D: Histogram of neighboring events detected by thresholding. The *x*-axis shows *Δx* of event *n* and the *y*-axis shows *Δx* of the following event, *n*+*1*. Panel E: Correlation of events detected by thresholding. Red indicates neighboring events occurring more frequently than random. Blue indicates neighboring events occurring less frequently than random. White areas indicate undefined correlation. Panel F: Histogram of neighboring events detected by MDL. Panel G: Correlation of events detected by MDL.

**Figure 5:**
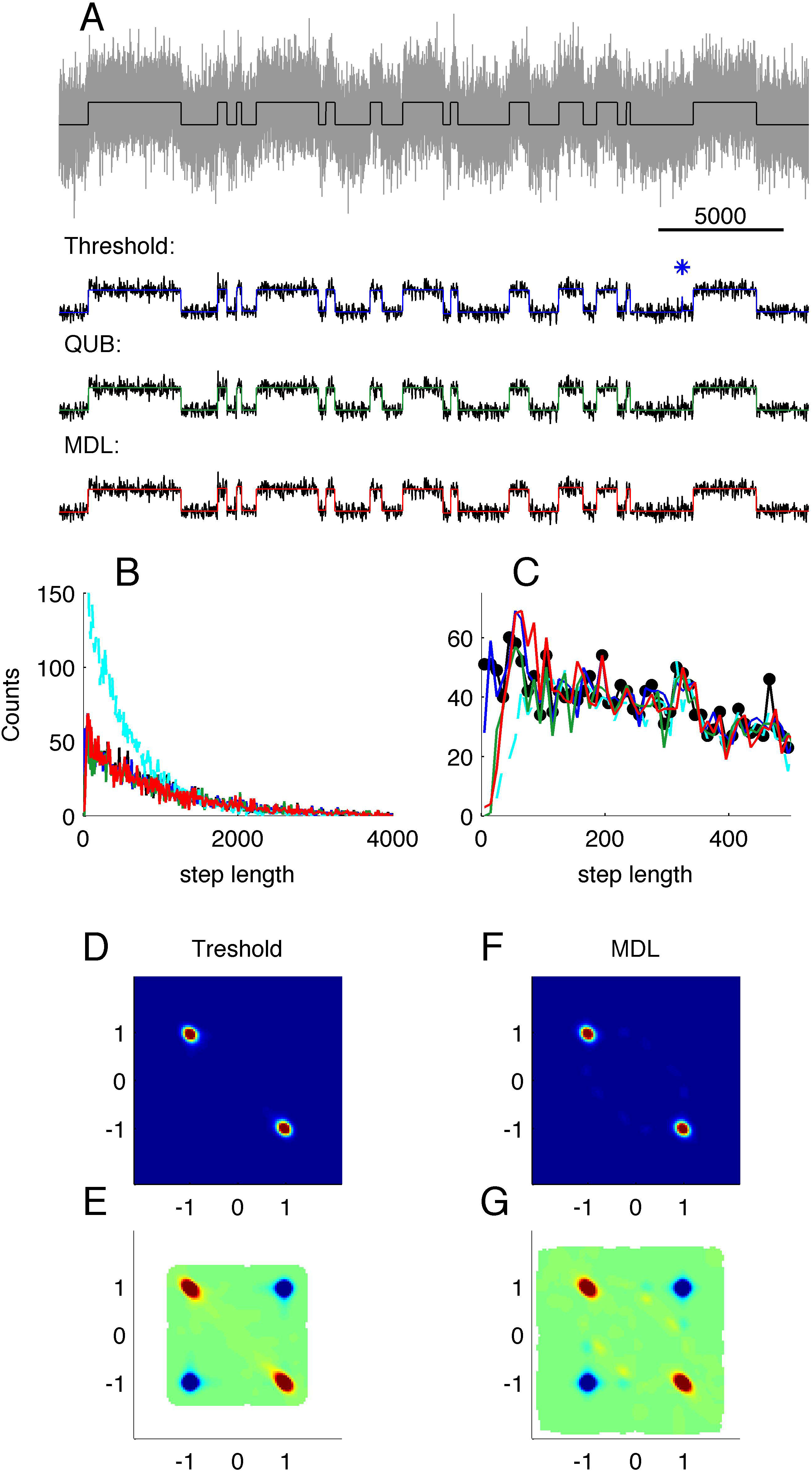
Validation on complex simulation data with different levels of noise. Data simulates three identical, but independently operating channels. First column, Panels A1-D1, is the case for low noise (SNR = 33); second column, Panels A2-D2, is for medium noise (SNR = 3.3); and last column, Panels A3-D3, is for high noise (SNR = 1). Panel A1-A3: Gray, segment of raw data (upper line only); black, true states; red, MDL idealization. Panels B1-B3: Histogram of detected steps. Panels C1-C3: 2D conditional histogram of neighboring steps. Panels D1-D3: Correlation between neighboring steps.

To compare our MDL-based detection methods to other methods (**Figure 4**), we used simulated test data consisting of *N = 5*10^6^* data points generated by a two-state Markov Model with equal probability for each state. Here, the transition probability of the Markov chain was 10^-3^ in each time-step and the standard deviation of the Gaussian noise was equal to the step size between Markov states, giving a SNR equal to 1. The slightly slower transitions kinetics used for test data in **Figure 4** compared to the test in **Figure 1** were chosen to provide test data for which the methods to be compared would all perform reasonably well.

We then applied MDL-detection to a simulated recording of three independent ion-channels with a complex structure of sub-conductance states (**Figure 5**). Each channel had one closed state (*C_0_*) and 4 open states (*O_1_, …, O_4_*) linked according to

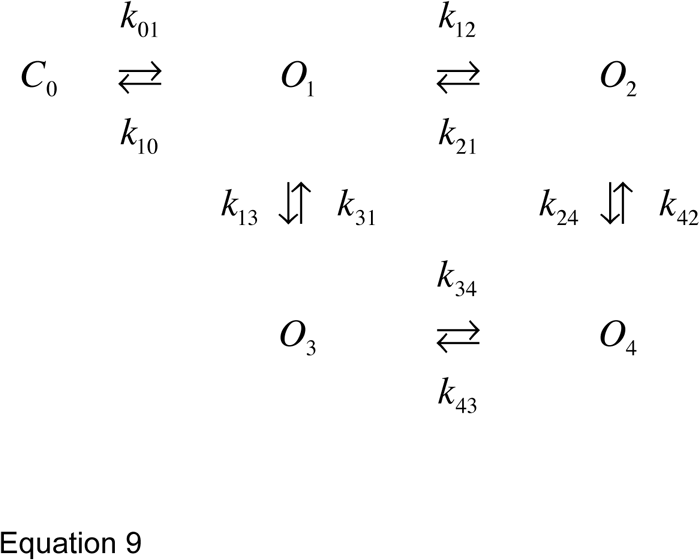

The amplitudes from the states *C_0_, O_1_,…, O_4_* are 0, 0.1, 0.3, 1, and 1. Thus, *O_3_* and *O_4_* are degenerate fully open states whereas *O_1_* and *O_2_* are nearly closed states but with some residual current. The rate constants are presented in Table 1. The rates of transition from state *O_2_* are relatively faster than the other rates, thus making *O_2_* a short-lived state.

**Table 1.**
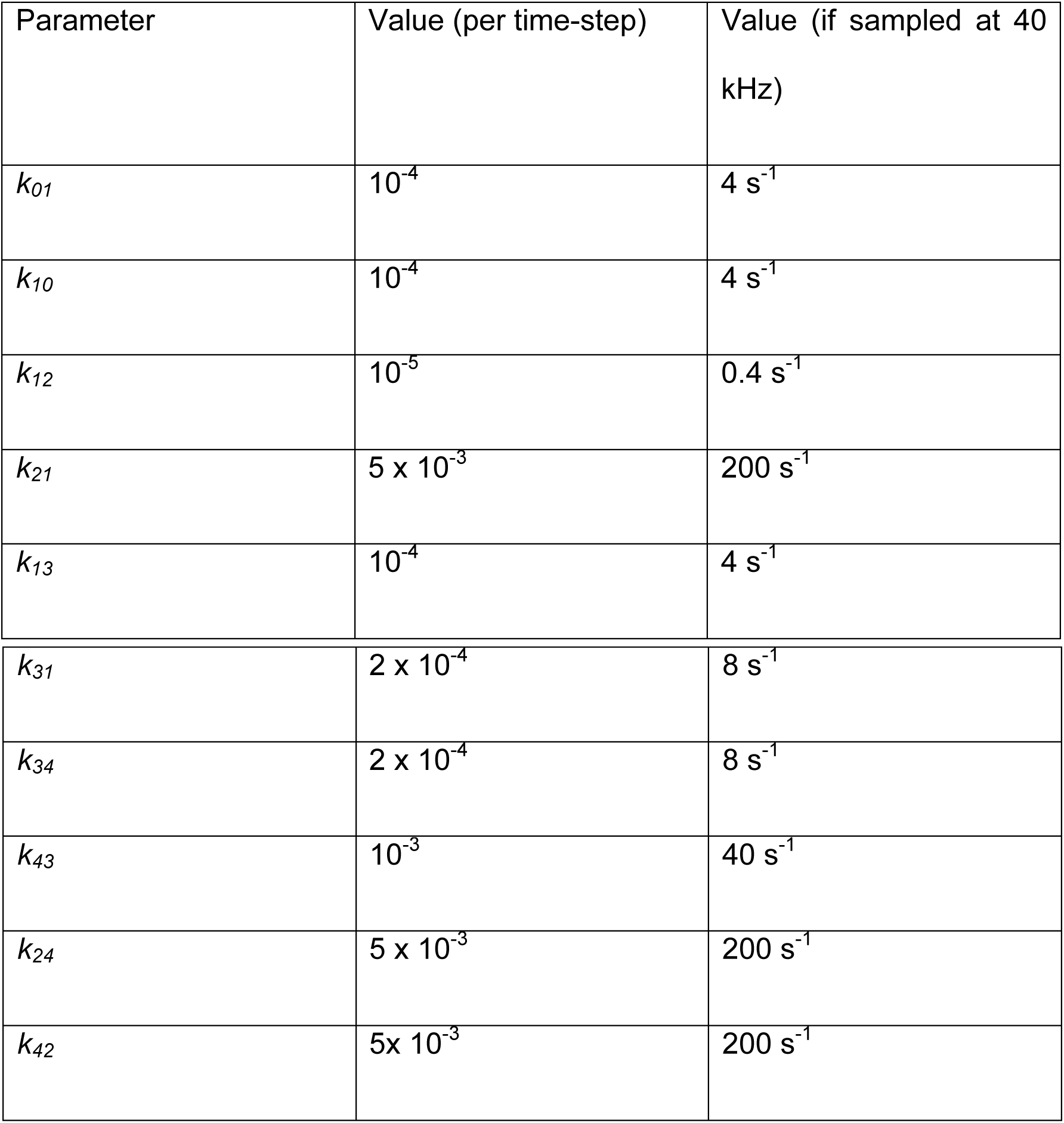
Transition kinetics of simulated complex channel used in **Figure 5**. States are described by Equation 9.

### Threshold-crossing and QuB analysis of synthetic data

We tested the MDL method against a threshold-crossing algorithm and the Viterbi algorithm as implemented in QuB (Rabiner, 1989; Nicolai and Sachs, 2013). The test was performed under optimal detection conditions for the competing algorithms: For the threshold crossing algorithm, we used prior information of the unitary steps in the process to set the detection threshold to 0.5. We applied low-pass filtering of the time-series in order to ensure reliability of the step detection by threshold crossing. In the next step we tested the sensitivity of the threshold crossing method against low-pass filtering by applying three different filters with cutoff frequency at 1/20, 1/40, and 1/80 times the Nyquist frequency (corresponding to 1, 0.5, and 0.25 kHz if the time-series modeled a real recording sampled at 40 kHz). Low-pass filtering was done using a digital three pole Butterworth filter. For easier comparison with MDL, each segment between detected steps was assigned its own mean value based on the unfiltered data points in the segment. The entire step-detection algorithm was coded in MATLAB.

For the sake of comparison, the same simulated data were also analyzed using the QuB software (Nicolai and Sachs, 2013). This approach uses the Viterbi algorithm to provide the most likely path through the state space of the HMM given the observed data. To provide the best possible conditions for event detection by this method we fitted data to the same two-state Markov model that was used to generate the test data.

### QuB analysis of hPIEZO1 channel unitary current

Data containing multiple open levels which varied in amplitude were first analyzed in QuB using the Segmental K-Means (SKM) algorithm (Qin et al., 1996b;1997) to idealize the events. We used a model with one or two open states to find the levels. The MDL algorithm was subsequently applied to the same dataset in order to evaluate its performance against the model-based approach. In the two-state model (SKM2), the initial estimates of the current in the closed and open states were assigned manually before idealizing the time series. In the three-state model (SKM3), we included a sub-state for which we set the starting estimate to 0.4 pA. Here, the three states were connected linearly and subjected to detailed balance. SKM recognized all occurrences of the predefined states from the entire trace; i.e. the two-state model recognized two states (closed and open) and the three-state model recognized the presence of three states (closed, low-conductance open-state and high-conductance open-state).

## Results

### Validation on simulated data

We first asked how well our method resolved a simple sequence consisting of a simulated single channel embedded in Gaussian noise. The channel was modeled as a two-state Markov process of unit step-size, with equal probability of being either open or closed, and with a state transition probability of 1/100, and the added Gaussian noise was set at SNR = 3.3 or SNR = 1 (**Figure 1**A top and lower panel respectively). On average, 98% of the steps were detected at SNR = 3.3 (low noise) and only 50% at SNR = 1 (high noise). The fraction of detected steps did not depend on record length (**Figure 1**B, solid green, low noise; solid blue high noise), whereas detection deteriorated in longer records when using the binary search alone, in particular at high noise levels (**Figure 1**B, dashed green, low noise; dashed blue high noise).

We next asked if the algorithm would generate false positive steps (**Figure 1**C). In general, false positives occur when a subset of consecutive data points randomly has a mean value sufficiently different from the rest of the segment to fulfill the MDL criterion. When analyzing any data set with many segments, there will be some finite probability that this occurs. To determine this probability, we constructed data series with a different total number of data points, *N*, but without deliberate introduction of any steps. For segments with *N* > 100, the method detected virtually no false positive steps. The probability of erroneously dividing the segment one or more times was 6 ± 1% for segments with 10 < *N* < 100 (**Figure 1**C, black). Only in segments with N < 10 was there any appreciable risk of detecting false positive steps. Increasing the minimum acceptable segment length, *l*_min_, to 3 or 5 reduced the risk of false positive steps in short segments, but had negligible effect on event detection in longer segments (**Figure 1**C; *l*_min_ = 1, black, *l*_min_ = 3, red, *l*_min_ = 5, blue).

Thus, the risk of generating false positive steps is greater within short segments (**Figure 1**C). This observation has implications for application of our method for analyzing data with fast kinetics under high SNR. For example, 63% of the segments of the input data in **Figure 1**A are shorter than 100 data points. Under conditions of high SNR, where the algorithm is able to detect most true transitions, some of the resulting segments can be sufficiently short to risk instances of false positive segmentation. In order to differentiate between genuine transitions and errors, we compared the expected precision (Equation 5) of the transitions (based on the length of adjacent segments) and the observed step (lmin = 1). Under high SNR, we found that the amplitude deviated more than expected on the basis of the precision estimate in 5% of the identified transitions, and that many of these occurrences had small |*Δx*| (**Figure 1**D, SNR = 3.3, dots indicate observed step and estimated precision. Red lines indicate 4 standard deviations from unity). In data with low SNR, the algorithm does not capture the shortest segments, and false positives will thus be rarer. Under low SNR, 4% of the transitions lay outside the expected range (**Figure 1**E, SNR = 1). This was presumably because the algorithm stopped before all segments were detected. In biological data, where we do not always have prior knowledge of typical step amplitudes and kinetics, we used *l_min_* = 3 to reduce putative false positive detections.

To verify that our method gave relevant information of the state-transitions, we calculated the joint probability histogram of neighboring steps. As the step generation mechanism was by definition a two-state Markov process (simulating a single simple channel with open and closed states), each step must necessarily be followed by an inverse step. We found that the conditional histogram showed exactly this predicted behavior: the vast majority of +1 steps (transition from closed to open state) were followed by −1 steps (transition from open to closed states) and vice versa (**Figure 1**F, low noise; **Figure 1**H, high noise). The correlation plot revealed the Markov structure of the process: +1 steps correlated positively with ‘-−1’ steps, and vice versa, whereas +1 steps correlated negatively with other +1 steps (**Figure 1**G, low SNR; **Figure 1**H, high SNR).

The above analysis showed that the algorithm resolved long segments, whereas sometimes missed short segments at low SNR. To characterize further the performance of our algorithm, we calculated the coherence spectrum between the input time series and the idealized output (**Figure 2**). Coherence was nearly 1 at low frequencies (indicated as inverse segment length) but decayed at high frequencies. At SNR = 1 (**Figure 2**, purple), coherence was ∼0.5 at frequencies corresponding to segments of length *n*∼100. This is consistent with our observation above that for a process with mean segment length 100 and SNR = 1, MDL detected roughly 50% of the segments (**Figure 1**B, solid blue). The observed roll-off at higher frequencies indicates that the idealized output failed to resolve shorter segments. At higher SNR the roll-off occurred at higher frequencies (higher values of 1/n), indicating that our algorithm was able to resolve increasingly finer details of the input (**Figure 2**; SNR 3.3, yellow; SNR 10, orange; SNR 33, blue). Thus, our algorithm resolves transient details, but in a manner limited by the SNR. It follows that our algorithm can resolve noisy data, even with SNR < 1, if the kinetics ensure that typical segments are sufficiently long.

### The combination of binary and tertiary search saves computation time

We determined the computation time for MDL to resolve data as shown in **Figure 1**A on a typical laptop computer (**Figure 3**). With SNR = 3.3 (as in **Figure 1**A upper), computation time scaled linearly with sequence length and was uniform across 10 trials. For example, the processing time for a sequence of 500,000 data points had a mean of 7.2 s (range 6.8 to 8.9 s) (**Figure 3**, blue circles show mean and error bars show range). With SNR = 1 (as in **Figure 1**A lower), the computation time was significantly higher and more variable between trials. For example, the mean processing time for 500,000 data points was ∼1000 s (range 75 – 3600 s) (**Figure 3**, red dots show mean and error bars show range from n = 10 iterations).

The computation time for uniform random data was even longer, did not vary between trials, and the processing time scaled with the square of the sequence length (**Figure 3**, yellow dots computation time for a single trial). To provide a comparison with HMM-based methods, we found the processing time for analyzing the sequences using our MATLAB implementation of the Viterbi algorithm to be linear and independent of SNR (**Figure 3**, purple asterisks show computation time for a single trial at SNR = 3.3).

The differences in computation time reflect which method is most involved in the search for breakpoints. While the binary search is fast, albeit apt to miss breakpoints in long segments, the tertiary search is computationally expensive, but is required to enable uniform recovery at different record lengths (**Figure 1**B, compare dashed and solid). However, our approach of combining binary and tertiary searches reduces computation time and enables computationally efficient searches on datasets with multiple breakpoints.

### Comparison with other idealization methods

Our motivation behind developing the MDL algorithm is to enable unbiased event detection and idealization of real electrophysiological data. The algorithm is designed to idealize ion channel recordings without user-specified inputs, and with the fewest possible assumptions, thus providing a versatile and general tool. We concede that other methods of idealization might be optimal for a particular system, thus providing more information in a particular application. In order to compare the fitness of other specialized methods to MDL, we analyzed a simulation consisting of a simple two-state Markov model emitting 0 or 1 with equal probability and Gaussian noise with standard deviation 1 (**Figure 4**A). The transition probability between states was set to 10^-3^ per time step, a 10 times slower process than used above (**Figure 1** - **Figure 3**). The distribution of step-lengths, i.e. the dwell time in each state, followed an exponential decay function (**Figure 4**B and C, black).

In this comparison of methods, we first applied threshold-based event detection on the simulated data. The threshold was set at 0.5, i.e. the 50% level between the two states. At the noise level used in the test, threshold-crossing required prior low-pass filtering of the data. For this, we used a 3- pole Butterworth low-pass filter, and tested the effect of three levels of low-pass filtering, i.e. at *F_Ny_/20, F_Ny_/40, and F_Ny_/80* (these filters correspond to 1, 0.5, and 0.25 kHz in relation to 40 kHz sampling).

The algorithm’s ability to detect short duration events using a threshold proved to be highly dependent on the filter width, with best sensitivity at *F_Ny_/40* (**Figure 4**A, dark blue, **Figure 4**B and C, dark blue – note that the dark blue overlaps other colors, with the exception of cyan in **Figure 4**B). With too little filtration, i.e. *F_Ny_/20*, the threshold algorithm detected many false-positive events (**Figure 4**B, cyan). On the other hand, with excessive filtration of the input data, i.e. *F_Ny_/80*, there was a penalty in the ability of the threshold-based algorithm to detect short segments (**Figure 4**C, cyan).

We next analyzed the data using QuB (**Figure 4**A, green). This approach uses the Viterbi algorithm, which provides the most likely sequence of states given an observed sequence of emissions, and provided that the probability of observing a particular emission from each state is known (Rabiner, 1989), without requiring low-pass filtering. The overall distribution of dwell times was similar to the input sequence (**Figure 4**B, green), although the probability of detecting transitions with a dwell time less than ∼30 time-steps was lower than for the case of threshold-based detection (**Figure 4**C, compare green and dark blue).

We then applied the MDL method to the data (**Figure 4**A, red). Overall, MDL performed well on the test data, and the distribution of dwell times matched the theoretical result (**Figure 4**B, red) (see also **Figure 2**, SNR = 1, typical segments in the input has *n*=1000 data points). The probability of detecting segments briefer than 30 time-steps was reduced to the same degree as seen with QuB. With MDL, there was a slight overrepresentation of segment lengths between 50 and 100 time steps, which we did not see with QuB. This bias of MDL is presumably due to the same effect as illustrated in **Figure 1**C, where brief segments face a certain risk of being erroneously sub-divided. The lower detection limits observed by QuB and MDL are determined by the SNR of the input data, rather than being an intrinsic property of these methods. The event detection by MDL had a similar distribution and correlation pattern as did the threshold crossing method (compare **Figure 4**D and E, with F and G).

These simulation results show that it is possible to optimize detection methods by taking into account prior knowledge of the process, such as kinetics and step size. The highest temporal resolution was found with the optimally filtered threshold-based method, and the accuracy of detection of true amplitudes was highest with the HMM based method. The MDL-based detection method, on the other hand, is designed to be generally applicable. When applied to simple processes it may provide an independent confirmation of the sorts of assumptions and constraints used in more refined analyses.

### Detection of sub-states in multiple channels

Many types of data do not lend themselves to analysis by conventional methods. For example, currents arising from background channels can be present in the data recording, or individual channels can exhibit transitions between sub-states. In these cases, MDL-based detection is still reliable. As a test of our algorithm on complex data of this type, we simulated the combined output of three simultaneously active, independent, and complex ion-channels, each as given by Equation 9. The simulation generated output from states with long and short dwell times and different current amplitudes. The expected amplitude steps defined in the dataset are ± 0.1 (from *C_0_* to *O_1_*), ± 0.2 (from *O_1_* to *O_2_*), ± 0.9 (from *O_1_* to *O_3_*) and ± 0.7 (from *O_2_* to *O_4_*). Transitions from *O_3_* to *O_4_* will not give a step in current, but these two states have different transition kinetics to the other states. We analyzed the same simulated dataset with different levels of noise (**Figure 5** A1-A3 shows part of the test data with noise (gray), original state emissions (black), MDL idealized (red)).

The first idealization of the recordings was in a condition of low noise (**Figure 5** A1, black and red traces superimposed, SNR = 33. The MDL algorithm detected multiple steps at the same amplitudes as defined for the channel (± 0.9, ± 0.7, ± 0.2, and ± 0.1; **Figure 5** B1, compare black and red). There was a small fraction of false-positive detections clustered about the origin of the histogram. Based on a comparison with **Figure 1**C, this bias is most likely due to the favored detection of very short segments under a condition of low noise.

The complex state transitions of the simulated channels (Equation 9) were apparent in the joint probability histogram of neighboring steps. This histogram depicted numerous open events followed by closing events (**Figure 5** C1, along the diagonal in the lower right quadrant). and also many closing events followed by open events, **Figure 5** C1, along the diagonal in the upper left quadrant). In addition, numerous off-diagonal events were observed.

We see a similar pattern in the corresponding amplitude correlation map (**Figure 5** D1). Here, the most highly correlated transition was +0.7, and the second most common -0.7, and vice versa. This result is doubtless because of the rapid flickering between *O_2_* and *O_4_*, as defined in the model. Events with a -0.1 step (transition from *O_1_* to *C_0_*) were highly correlated with a +0.1 step (transition from *C_0_* to *O_1_*). We also found steps of -0.7 to be highly correlated with steps of +0.7 (**Figure 5** D1, upper left quadrant, reflecting transitions from *_O4_* →*O_2_* →*O_4_*) and also with -0.2 transitions (reflecting *O_4_* → *O_2_* → *O_1_*), and note that the *O_4_*→*O_2_* transition seemed more strongly correlated, whereas the *O_3_*→*O_1_* transition was less correlated. These results presumably arise from the kinetics of transition away from *O_1_* being slower than from *O_4_*. Therefore, in the presence of other channels, slow transitions like *O_3_*→*O_1_*→*O_3_* are not often observed directly in step detection, because they are likely to be interrupted by events in other channels.

We next analyzed the same simulated recording under increased noise levels (**Figure 5** A2, SNR = 3.3; and A3, SNR = 1). As expected, the sensitivity of detection gradually declined for small amplitude events, and the estimation of the plateau magnitudes became more influenced by the SNR (Compare red and black lines in lower panels of **Figure 5** A2 and A3 and in B2 and B3). The joint probability histogram and correlations between neighboring steps were remarkably robust, and most transitions observed under very low noise (**Figure 5**C1 and D1) were also detected in recordings with moderate noise (**Figure 5**C2 and D2). At high noise, only the most dominant amplitude events were discernable. In that condition, it was no longer possible to clearly distinguish ± 0.7 from ± 0.9 events. Even though the detection limit was limited by noise, it remained possible to identify *C_0_*→*O_1_*→*O_3_* (**Figure 5** D3, small peak at [Δx_n_, Δx_n+1_] = [0.1, 0.9]), *O_1_*→*C_0_*→*O_1_* (**Figure 5** D3, small peak at [Δx_n_, Δx_n+1_] = [-0.1, 0.1]), and *O_3_*→*O_1_*→*C_0_* (**Figure 5** D3, small peak at [Δx_n_, Δx_n+1_] = [-0.9, -0.1]).

Thus, our analysis of simulated recordings predicts that the algorithm should successfully resolve subconductance structures even under high noise conditions and in the presence of potentially interfering signals arising from multiple channels.

### Analysis of human PIEZO channels

We tested the MDL method for analysis of experimental data from human PIEZO1 ion channels. The PIEZO1 channel is a mechanosensitive receptor that is activated by membrane tension (Cox et al., 2016), which can be applied experimentally by stretching patched membranes with either positive or negative pressure (Besch et al., 2002). This channel activates and deactivates rapidly, tracking closely the onset and termination of the pressure stimulus. It also shows voltage-dependent inactivation, which is faster at hyperpolarized membrane potentials. In our preparation we observed openings to multiple levels during application of steady pressure; some of these open levels may in fact represent sub-conductance states of the channel, affording a useful test of the performance of our MDL algorithm.

The analyzed segment lasted 140 s and contained 14 episodes of applied stimulus, each of 5 s duration, with a 5 s relaxation interval between the stimuli. We held the patch at five different potentials during the recording (**Figure 6**A shows the voltages and **Figure 6**B the suction pulse train that served as the mechanical stimulus; **Figure 6**C shows the recorded current). The drift in the baseline leakage current resulting from changes in voltage was cancelled using the baseline algorithms in QuB (www.qub.buffalo.edu), whereas the MDL method was applied to the unfiltered current trace. MDL detected the channel opening events even though the unitary current amplitude changed as a function of the potential applied across the patch (**Figure 6**C, compare amplitude of steps in red, blue, and green highlights). The MDL algorithm also detected current steps of multiple sizes (**Figure 6**C, putative sub-state currents indicated by arrows in red and green highlights). If the membrane potential was -60 mV or more negative, the current steps appeared to reflect openings of different duration and current (**Figure 6**C, arrows in red and green highlight indicate small current events). At low polarization (-20 mV), these events were detected only rarely, presumably because their amplitude fell below the noise detection limit. The algorithm detected 125, 191, 985, 230, and 632 events at -20 mV, -40 mV, -60 mV, -80 mV and -100 mV respectively. A total of 78 events that were detected in proximity to recording artifacts occurring at changes in holding potential were ignored in the analysis (see asterisks in **Figure 6**C).

**Figure 6:**
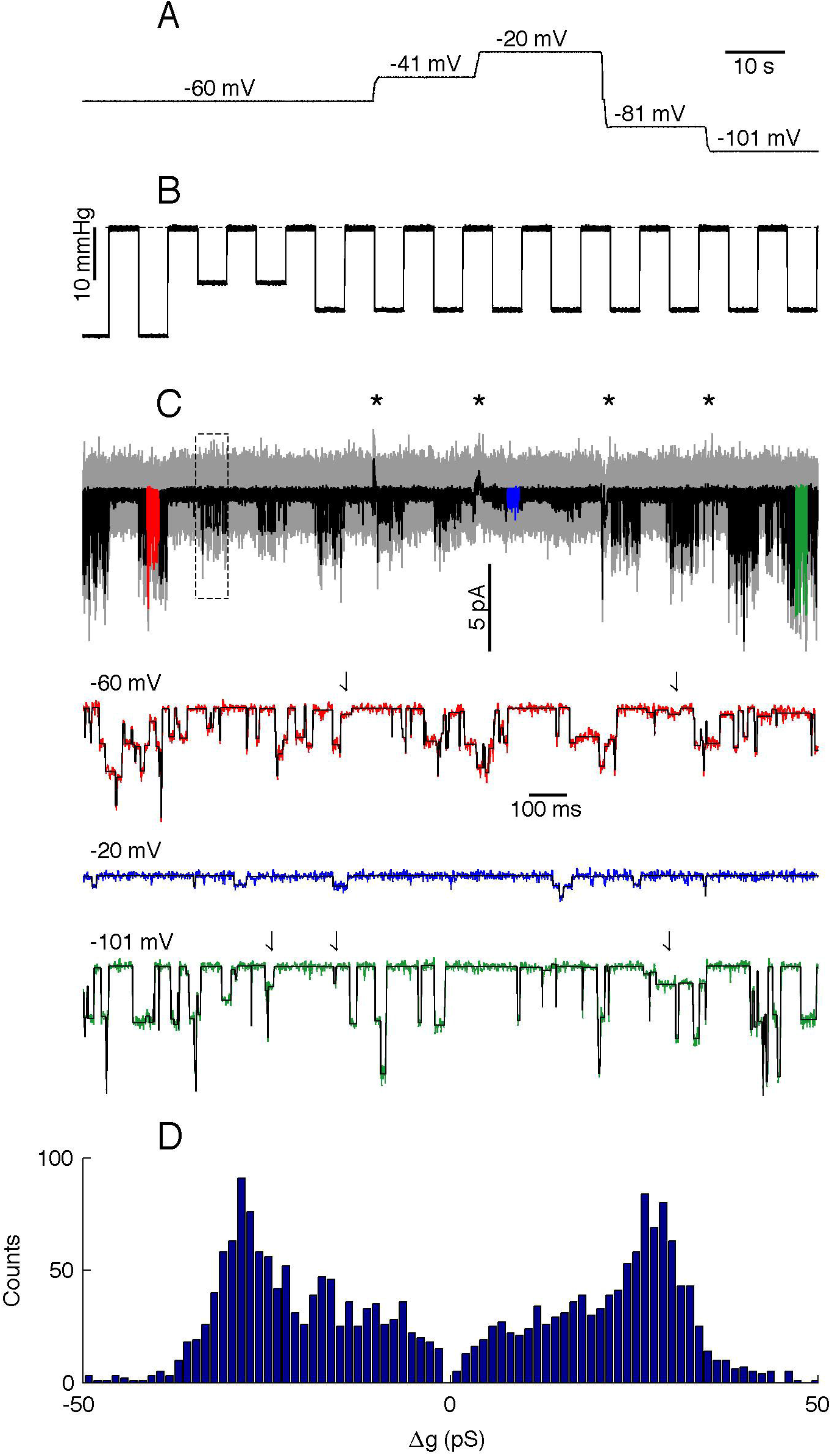
Analysis of PIEZO1 channels. Panel A: Membrane potential during the recording. Panel B: Mechanical stimulus, dashed line indicates 0 mmHg. Panel C: Analyzed current trace. Gray is unfiltered raw data; black is low-pass filtered data. Black asterisks indicate capacity artifacts during changes in membrane potential; the rectangle indicates the region analyzed in Figure 8. Colored segments (red, blue, and green) are shown in detail below with the idealized trace in black. Arrows in highlighted traces (red and green only) indicate putative sub-state currents. Panel D: Histogram of channel events.

To facilitate analysis across the different voltages of the time series, we divided the MDL-detected current steps by the concurrent holding potential to calculate changes in conductance, *Δg*. assuming channel conductance is linear. The event amplitude histogram presented a wide range of conductance steps with peaks at *Δg* = ± 28 pS, but we also observed smaller transitions down to *Δg* = ± 6 pS (**Figure 6**D). The relative fraction of smaller conductance steps varied with the holding potential. Thus, at -20 and -40 mV the algorithm detected relatively few steps with amplitude lower than 28 pS. However, higher holding potentials revealed a wide distribution of different amplitudes, although steps at ± 28 pS amplitude were always present and appeared to be the dominant amplitude.

We then analyzed the joint probability distribution and correlation of neighboring steps at different holding potentials (**Figure 7**A and B shows -60 mV, C and D shows -100 mV). The main transitions at ± 28 pS were clearly identified at all holding potentials (**Figure 7**A and C shows -60 and -100 mV, other holding potentials not shown). However, there were some notable differences as a function of the voltage. For example, at -60 mV the +6 pS open steps were frequently followed by a -6 pS closing step, whereas other sub-states were infrequent. However, at -100 mV holding potential there was a wider range of sub-amplitude openings, including ± 10 and ± 16 pS.

**Figure 7:**
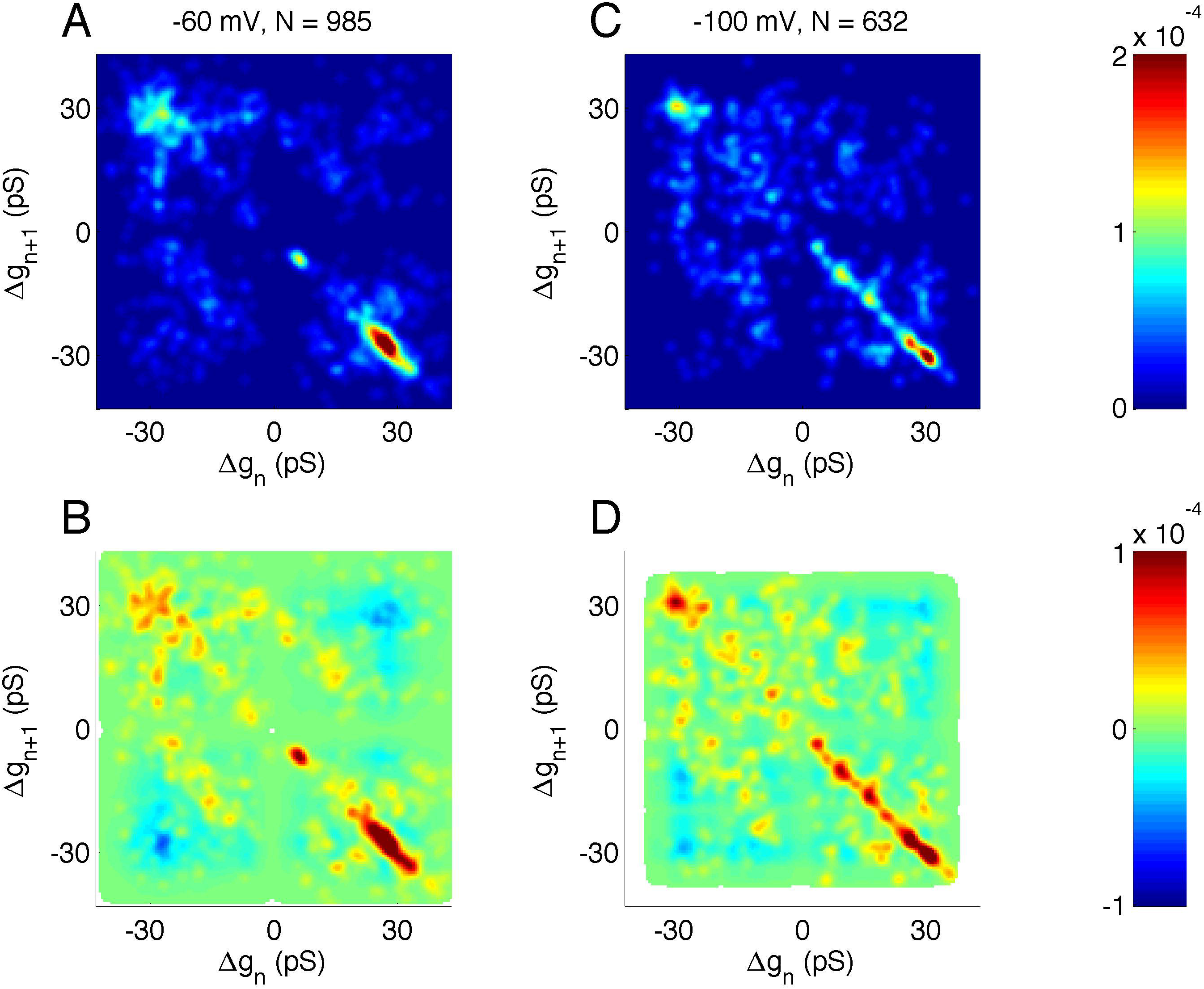
Correlation analysis of neighboring events in PIEZO1 channels recorded at -60 and -100 mV holding potentials. As in other figures, *x* and *y* scales are arranged so that open events followed by closed events populate the lower right quadrant. Panel A: Distribution of neighboring events at -60 mV. The *x*-axis shows *Δg* of event *n* and the *y*-axis shows *Δg* of the following event, *n*+*1*. Panel B: Correlation of neighboring events at -60 mV. Red indicates neighboring events occurring more frequently than random. Blue indicates neighboring events occurring less frequently than random. White areas indicate undefined correlations. Panel C and D: same as A and B but for -100 mV holding potential.

Multiple factors must account for the details of the observed channel kinetics, and a full characterization of hPIEZO1 channel properties is beyond the scope of this report. For the present, we emphasize that the unsupervised MDL idealization and our joint probability and amplitude correlation method proved to be sensitive to the changes in the full open amplitudes and sub-states. Thus, our method may provide a firm foundation for subsequent analysis using this model approach.

Several methods are already available for the analysis of single-channel data. To get an understanding of performance of the MDL method in comparison to the established techniques, we analyzed a segment of the hPIEZO1 data with QuB using the Segmental K-means (SKM) method to estimate the unitary channel amplitudes. Unlike MDL, this method explicitly employs a defined 2- or 3-state model, with user-defined rates emulating the inverses of the observed closed and open state lifetimes. We chose an arbitrary part of the recorded segment for the MDL analysis (**Figure 8**A, indicated by the rectangular inset in **Figure 6**C). The segment was recorded at -60 mV holding potential, where ± 6 and ± 28 pS were the dominant transitions according to the analysis above (**Figure 7** A and B).

**Figure 8:**
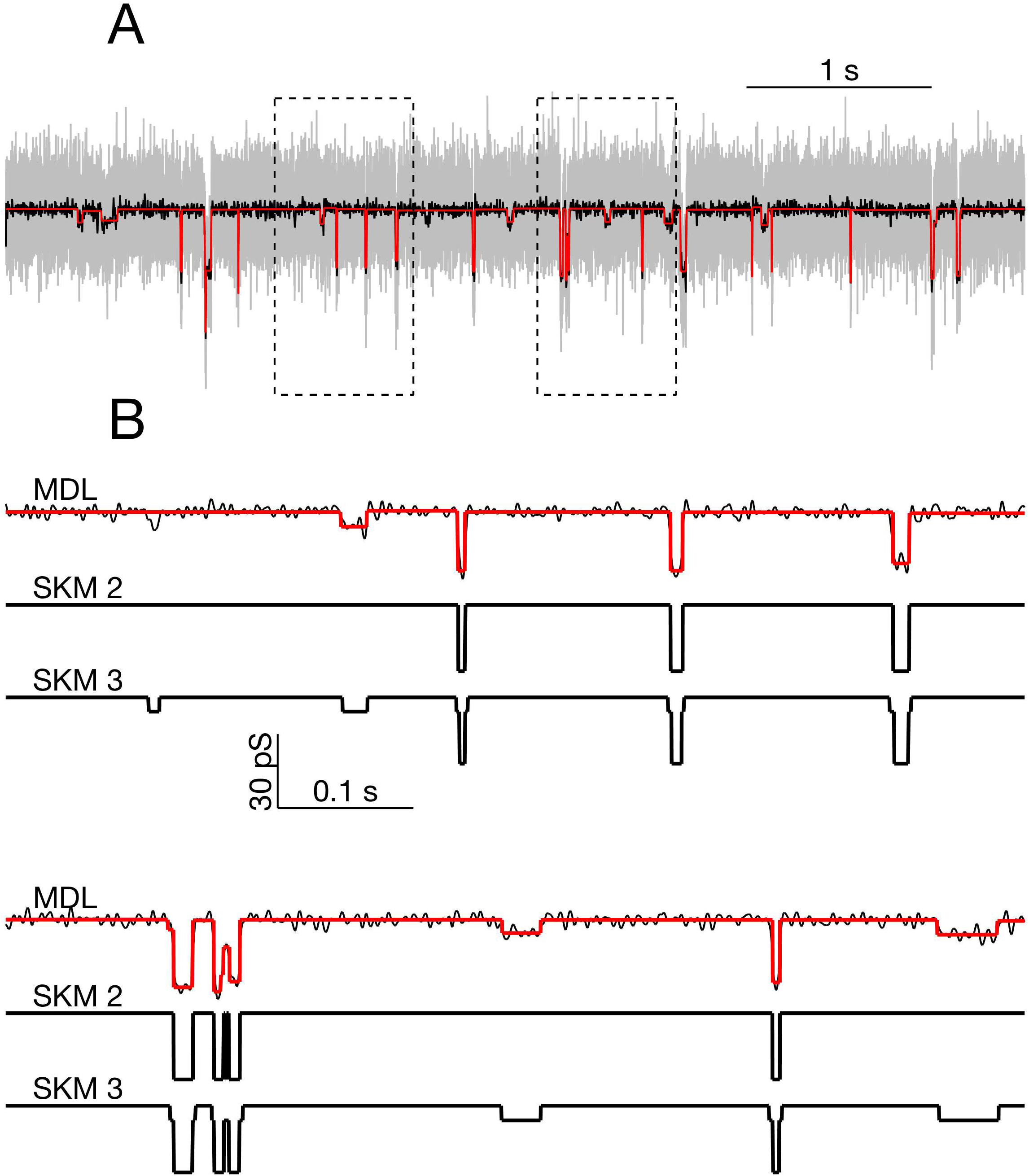
Comparison of MDL and SKM analysis on data from PIEZO1 channels recorded at -60 mV. Panel A: Analyzed segment (part of recording in Figure 6, indicated by rectangular inset in Figure 6C). Raw data, gray; low-pass filtered, black. MDL idealized, red. Rectangles indicate regions expanded below. Panel B: Expanded view of parts of the trace in panel A. Top: MDL idealized (red) and low-pass filtered data (black). Middle: Idealized by two state SKM (SKM2). Bottom: Idealized by 3 state SKM (SKM3).

We first idealized the segment using SKM for a 2-state ion channel. The idealized current trace contained 40 open or closed events. The analysis yielded a transition between the closed state and a single open state of 29.8 pS conductance. Visual inspection confirmed that the idealization detected the high amplitude openings but missed a number of smaller amplitude openings (Figure 8B, compare SKM 2 and SKM 3 or MDL).

When the segment was reanalyzed with SKM using a 3-state linearly-connected model, the idealized data contained 86 events. Now the event sizes were either *Δg* = ± 6.7 pS or ± 23.2 pS. The current at the fully opened state was 29.8 pS, equal to the sum of the two main event sizes. By visual inspection, the smaller event size appeared to reflect opening to the same sub-conductance state discussed above (see arrows in highlighted parts of **Figure 6**C).

We then idealized the data segment using the MDL method, which detected 61 events and proved to be sensitive to most of the visually-observed openings to both the sub-state and the fully-open state. The apparent amplitude of the events was roughly the same as for the 3-state SKM approach, although MDL appeared slightly conservative, in failing to capture some possible small amplitude events detected by the 3-state model (**Figure 8**B).

## Discussion

Analysis of currents from a membrane patch containing an unknown number of channels in the presence of a noisy background or leakage current is challenging, regardless of the applied algorithm. The various available methods all have different strengths and weaknesses. Most of the currently used methods require some degree of user-dependent input and supervision, through selection of a low-pass filter followed by application of an event-detection threshold, or a *priori* Markov models. While such methods can have good performance in analyzing records with a few simple channels, there is currently no way of accommodating complicated records, which may contain events of differing amplitudes and kinetics. While the HMM based methods use a kinetic model to estimate the individual channel current amplitudes, the observed amplitudes are relatively insensitive to the magnitude of the transition rate constants (Qin et al., 1996a). For dealing better with multichannel currents from identical channels, the MAC (macroscopic) algorithm in QuB (Milescu et al., 2005) is robust for estimating transition kinetics, the number of active channels in the pool, and the mean jump size arising from an arbitrary stimulus (Bae et al., 2013).

Here we show that the Minimum Description Length principle (MDL) enables idealization of patch recordings in a user-independent manner. We find that this method can be applied to raw data without the need to correct baselines, and has good performance even in cases where event amplitude changes during the recording (**Figure 6** and **Figure 7**). We provide proof of this principle based on three main findings. First, we applied our method to simulated data and investigated the limits for detection. These simulations showed our method to be efficient for detecting events with multiple independent Markov processes under various noise levels. Finally, we used a novel correlation analysis to relate the results to the transitions of single channels.

Our approach required the solution of two sub-problems: First, we needed an objective criterion for model selection. More precisely, it was necessary to test whether a complex model of the data using many discrete segments is superior to a simple model with fewer segments. For this type of assessment, several measures of quality have been proposed, among which the Akaike Information Criterion (Akaike, 1973), Bayes Inference Criterion (Schwarz, 1978), and the Minimum Description Length Principle (Rissanen, 1978;Lee, 2001). In our hands, MDL proved to work best, and we derived a cost function
that described the balance between model complexity and fitness. Optimizing the cost function posed a second challenge: Calculation of the description length depends on multiple interdependent variables, all of which contribute to the cost function. A full search of all possible combinations in solution-space is not universally applicable, since the expected number of events in real data is often in the thousands, which would be computationally impossible. Instead, we resorted to an iterative procedure in which breakpoints were inserted sequentially. One method for this is presented by the binary segmentation process (Scott and Knott, 1974;Kalafut and Visscher, 2008). However, when a binary search was applied to simulated data, the fraction of detected steps declined as a function of record length. In contrast, no such dependence was observed when implementing a tertiary search, where the algorithm searches for possible combinations of two break points before terminating (compare solid to dashed lines in Figure 1B). A binary search was used in the method presented earlier by Kalafut and Visscher (2008) for analyzing movement of the molecular motor kinesin. Presumably, loss of detection in long time series is less of a concern in their application because the motion of kinesin has a preferred direction, which makes it simpler to decompose the time series into segments. For ion channel applications, time series are often generated by a *stationary* random process, which makes it more unlikely that breakpoints inserted one by one fulfills the criterion given by Equation 4. Thus, resolution of typical data for our application also requires the more elaborate tertiary search method where breakpoints are inserted two by two. However, to optimize the computing time and reliability, our algorithm combines binary and tertiary searches (Figure 3).

We anticipate that there could be cases for which the tertiary search method will also be sub-optimal. For example, a long time-series constructed of alternating segments of *equal* length would require a higher SNR in order to be resolved than would be necessary if the segments were of *random* length, as in our tests. However, this situation is unlikely to occur in real data from ion channels for which stochastic processes drive the changes in conductance states. Furthermore, were such a circumstance actually encountered, there are convenient analysis methods for accommodating periodicity (Little et al., 2011).

Proper use of prior knowledge of the underlying biophysical processes generally enhances data analysis. For example, supervised methods such as SCAN take into account the presence of low-pass filters, and are thus able to determine transition points with accuracy exceeding the sampling rate of the data (Colquhoun and Sakmann, 1985). We found that the MDL method approached (but did not exceed) the performance of methods that explicitly use prior information (Figure 4). In situations where analysis can be performed by more specialized methods, the MDL-based method may nevertheless serve as an independent test of the model assumptions, and guide development of more refined analysis. Due to the minimal number of assumptions, our unsupervised MDL method may prove particularly useful for complex data from recordings of channels with unknown sub-state structure, and in cases where multiple independent channels are active (Figure 5 and Figure 8).

We applied the MDL method to experimental recordings of human PIEZO1 (hPIEZO1) channels, in which we encountered currents with multiple amplitudes (Figure 6 and Figure 7). Our analysis showed that the algorithm has particular advantage in situations where prior knowledge of the sub-conductance states of a channel is lacking. The flexibility of the MDL algorithm is a desirable property when performing analyses on large datasets. The MDL algorithm is generalized and can be used with time-series datasets acquired from other disparate sub-fields of biology characterized by state models (Nicolai and Sachs, 2014).

## Conclusion

We developed an idealization method based on the minimal description length principle for the use of analyzing ion channel recordings. The method was validated on simulated data, with characterization of event detection reliability under different noise and recording conditions. We the applied the algorithm to the analysis of patch clamp recordings of currents from the human PIEZO1 channel. Results of this test confirmed the algorithm’s fitness to detect sub-conductance states.

## Acknowledgement

This study was supported by the Lundbeck Foundation and the Dynamical Systems Interdisciplinary Network, University of Copenhagen (JKD), the Novo Nordisk Foundation, and the Danish National Research Foundation (JPH, MSN) and the NIH (FS, RS) R01HL054887. We note professional editing of the manuscript by Inglewood Biomedical Editing. MATLAB version of the MDL-algorithm is available at the File Exchange at MATLAB Central (Dreyer, 2016), and the algorithm is also available as a plug-in for the ion-channel analysis program QuB (Nicolai and Sachs, 2013).

This manuscript is submitted to Frontiers Neuroinformatics.

